# Basement membrane patterning by spatial deployment of a secretion-regulating protease

**DOI:** 10.1101/2024.07.06.602330

**Authors:** Hui-Yu Ku, David Bilder

## Abstract

While paradigms for patterning of cell fates in development are well-established, paradigms for patterning morphogenesis, particularly when organ shape is influenced by the extracellular matrix (ECM), are less so. Morphogenesis of the Drosophila egg chamber (follicle) depends on anterior-posterior distribution of basement membrane (BM) components such as Collagen IV (Col4), whose symmetric gradient creates tissue mechanical properties that specify the degree of elongation. Here we show that the gradient is not regulated by Col4 transcription but instead relies on post-transcriptional mechanisms. The metalloprotease ADAMTS-A, expressed in a gradient inverse to that of Col4, limits Col4 deposition in the follicle center and manipulation of its levels can cause either organ hyper- or hypo-elongation. We present evidence that ADAMTS-A acts within the secretory pathway, rather than extracellularly, to limit Col4 incorporation into the BM. High levels of ADAMTS-A in follicle termini are normally dispensable but suppress Col4 incorporation when transcription is elevated. Our data show how an organ can employ patterned expression of ECM proteases with intracellular as well as extracellular activity to specify BM properties that control shape.

## Introduction

Animal development requires generating spatial patterns within tissues in order to create a functional organ or organism. Decades of research has led to an understanding of general mechanisms that diversify cell fates along various body axes. For instance, morphogens secreted across a field of cells can create differential transcriptional states within cells in relation to their distance from a localized source (1), while lateral inhibition can specify discrete identities amongst initially equivalent neighboring cells (2). The process of morphogenesis ---the acquisition of physical form--is of equal importance to cell fate decisions in development. However, fewer paradigms exist for how spatial information is translated into tissue shape.

Shape ultimately involves regulating mechanical properties of not only cells but also non-cellular elements such as the extracellular matrix (ECM) (3–8). Spatial differences can be imposed through transcription, translation and also post-translational means that are unique to ECM’s extracellular nature. Such means include specialized secretory pathways to exit the cell, incorporation into pre-existing ECM, and protein processing and modifications that alter physical ECM properties. The list of processes where spatial differences in ECM drive organ shape is rapidly growing; recent examples range from axis extension of the early mouse embryo to folds in the Drosophila wing disc to branching of mammalian mammary glands (9–12). This new appreciation emphasizes the importance of uncovering mechanisms that generate ECM patterns across a tissue.

A powerful system to investigate ECM patterning has emerged in the Drosophila follicle, also called the ‘egg chamber’. This simple tissue begins development as spherical but then elongates along its anterior-posterior (A-P) axis during growth to form an ovoid egg (**Fig. 1A**); precise shape of this egg is required for full fertility (13, 14). Data have revealed that this morphogenetic change relies heavily on patterning of the ECM that surrounds the follicle (3–5, 7, 15, 16). The follicle ECM is a basement membrane (BM) that is in many ways typical of BMs that underlie most animal organs. BM components synthesized and secreted by follicle epithelial cells are assembled to create a stiffness gradient with mirror symmetry along the A-P axis, in which the organ center stiffens compared to the terminal regions that are more compliant. This ‘mechanical corset’ of relative stiffness alters morphogenetic cell behaviors to direct A-P rather than isotropic growth, thus guiding a precise degree of tissue elongation. Related phenomena generate BM mechanical differences during mammalian morphogenesis (9, 12), suggesting that mechanisms learned from the fly follicle may illuminate general principles that create organ-sculpting ECM.

**Figure 1.**
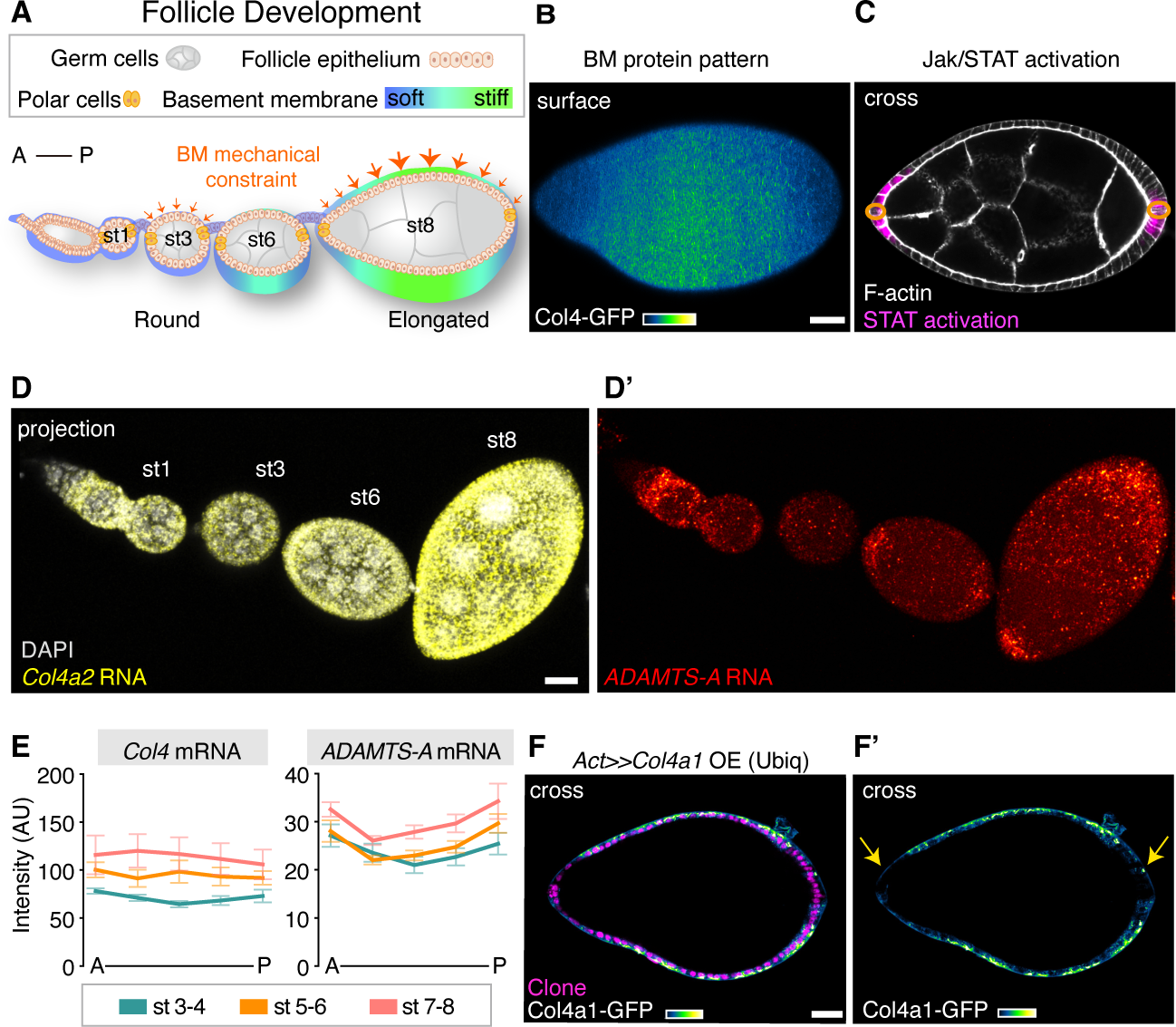
*ADAMTS-A* but not *Col4* shows an A-P gradient of transcription. (**A**) Overview of follicle morphogenesis from early-to-mid stages. Germ cells are covered by epithelia and two pairs of polar cells, with an underlying BM. The BM of young spherical follicles (st 1-3) is uniformly soft, subjecting the organ to homogenous mechanical constraints. As the follicle develops (st 5-8), a gradient of stiffer BM in the center shifts the constraints, leading to tissue elongation along the A-P axis. (**B**) BM pattern in stage 8 follicle (single plane, flattened). A heat map (blue to yellow: low to high) of endogenously tagged Col4-GFP shows intensity highest at center and tapering off toward termini. (**C**) STAT activation in follicle termini (magenta) mediated by Upd from polar cells (circled in orange), shown in cross-section. (**D**) HCR detects transcription of *Col4a2* (yellow, in D) and *ADAMTS-A* (red, in D’) in follicles). DAPI signal included for staging. Images are maximum projections. (**E**) Quantitation of Col4 and ADAMTS-A RNA distribution in stages 3 to 8, along five equal A-P divisions. Data plotted are mean ± SEM. Col4 expression is homogenous along A-P axis, with levels increasing during development. *ADAMTS-A* shows elevated expression at follicle termini as well as clear central expression. (**F**) Uniform overexpression (clone shown by magenta) of *Col4a1-GFP* transcript throughout the follicle leads to differential incorporation of Col4a1-GFP protein in center compared to termini (arrows). Cross-sections are shown. Scale bar: 20 μm.

A major feature of follicle BM patterning is a symmetric gradient of Collagen IV (Col4) levels that are high in the center and lower at the anterior and posterior poles (**Fig. 1A, B**) (17, 18). These levels roughly mirror the measured mechanical stiffness gradient. Col4 is present not only in a conventional planar matrix, but also in small fibril-like structures that confer distinct mechanical properties and are organized supracellularly (19–21). The gradient of overall Col4 levels is shared by a graded density of these fibrils (13, 21). Both are required for the tissue material properties that confer egg shape.

What mechanisms dictate the specific BM pattern that determines the fecundity of the animal? Formation of Col4 fibrils requires a specialized secretory pathway, and their orientation and supracellular structure depends on a collective epithelial migration (19, 20). We showed recently that a Drosophila matrix metalloproteinase (MMP1) is secreted from a pair of cells at anterior pole of the organ (‘polar cells’) and acts extracellularly to finely tune the stiffness gradient through promoting fibril formation at the follicle anterior (13). However, neither follicle cell migration nor MMP1 impact the overall levels of Col4. Here we identify an unexpected mechanism that regulates follicle BM patterning, involving intracellular control of Col4 secretion by a disintegrin-like and metalloproteinase with thrombospondin type 1 motif protein ADAMTS-A.

## Results

### The follicle Col4 pattern is not controlled by transcription

The Col4 gradient of stage 6-8 follicles displays levels lower at the terminal regions than at the center (**Fig. 1B**). Terminal versus central patterning of cell fates in the epithelium is controlled by the morphogen-like cytokine Upd, which is secreted from polar cells to activate the JAK/STAT pathway in nearby epithelial cells (**Fig. 1C**) (22). Disrupting Upd signaling homogenizes Col4 levels and alters follicle shape (17, 23), but the mechanisms through which follicle BM pattern is regulated remain unclear.

Since the Col4 gradient is inverse to that of STAT activity, one simple mechanism to generate the gradient would be if Upd signaling regulates Col4-encoding gene transcription. We visualized transcripts of *Col4a2* using hybridization chain reaction (HCR) and saw no differences in levels along the follicle AP axis before st. 8; this is consistent with in situ hybridization, expression of enhancer traps in Col4 regulatory regions and available scSeq data from follicle epithelia (24)(**Fig. 1D-E, S1A, B**). We also repeated experiments that overexpress Col4 from heterologous promoters (17). Even when a Col4-encoding transcript is expressed at uniformly high levels via follicle-wide induction of the *Actin-GAL4*-driven FLP-out system (25) (hereafter ‘ubiquitous FLP-out’), protein expression is severely reduced at the follicle termini (**Fig. 1F**). Therefore, post-transcriptional mechanisms must shape the Col4 gradient.

### The central region of an ADAMTS-A gradient is required for tissue mechanics and shape

Metalloproteases are well-known post-translational regulators of ECM. Drosophila MMP1 influences BM fibril formation and therefore egg shape, but it does not affect overall Col4 levels and this pathway acts independently of STAT signaling, excluding MMP1 as a gradient regulator (13). In a screen for transcriptional targets of STAT in the follicle, Wittes and Schüpbach identified the metalloprotease ADAMTS-A and found that it is also required for elongation (26). *ADAMTS-A* is elevated in follicle termini, leading to a simple model that ADAMTS-A may locally degrade BM components to soften the terminal regions. We reexamined *ADAMTS-A* expression using HCR and confirmed terminal elevation in stage 6-8 follicles, but also noted clear though lower levels throughout the follicle center. *ADAMTS-A* was also detected at uniform levels during earlier stages 2-5 (**Fig. 1D-E, S1A**). The data document varying levels of *ADAMTS-A* along the A-P axis, in a pattern reciprocal to the Col4 gradient.

We first explored ADAMTS-A function in follicle shape by tissue-wide depletion and overexpression. Tissue-wide depletion using ubiquitous FLP-out resulted in hyperelongation (**Fig. 2A-B**). This is distinct from the rounding seen by Wittes and Schüpbach using *GR1-Gal4*, perhaps due to weak early expression of *GR1-GAL4* (27, 28); in ADAMTS-A depletion by *tj-GAL4*, fusion between early follicles could be seen, complicating analysis. Tissue-wide overexpression with ubiquitous FLP-out, *GR1-GAL4* or *tj-GAL4* all resulted in round follicles, a phenotype opposite to that seen when ubiquitous FLP-out was used for depletion (**Fig. 2C**). When ADAMTS-A overexpression was restricted to the follicle center initiating at st. 6 via *mirr-GAL4* (17, 29), it caused strong rounding, a phenotype opposite to the hyperelongation seen when ADAMTS-A is depleted from the same region (**Fig. 2D-G**). Wittes and Schüpbach further manipulated ADAMTS-A using a *fru-GAL4* elevated in terminal regions, but because this driver shows uniform expression in young follicles and would deplete the central domain of ADAMTS-A as well (**Fig. S1C**), we chose not to use it.

**Figure 2.**
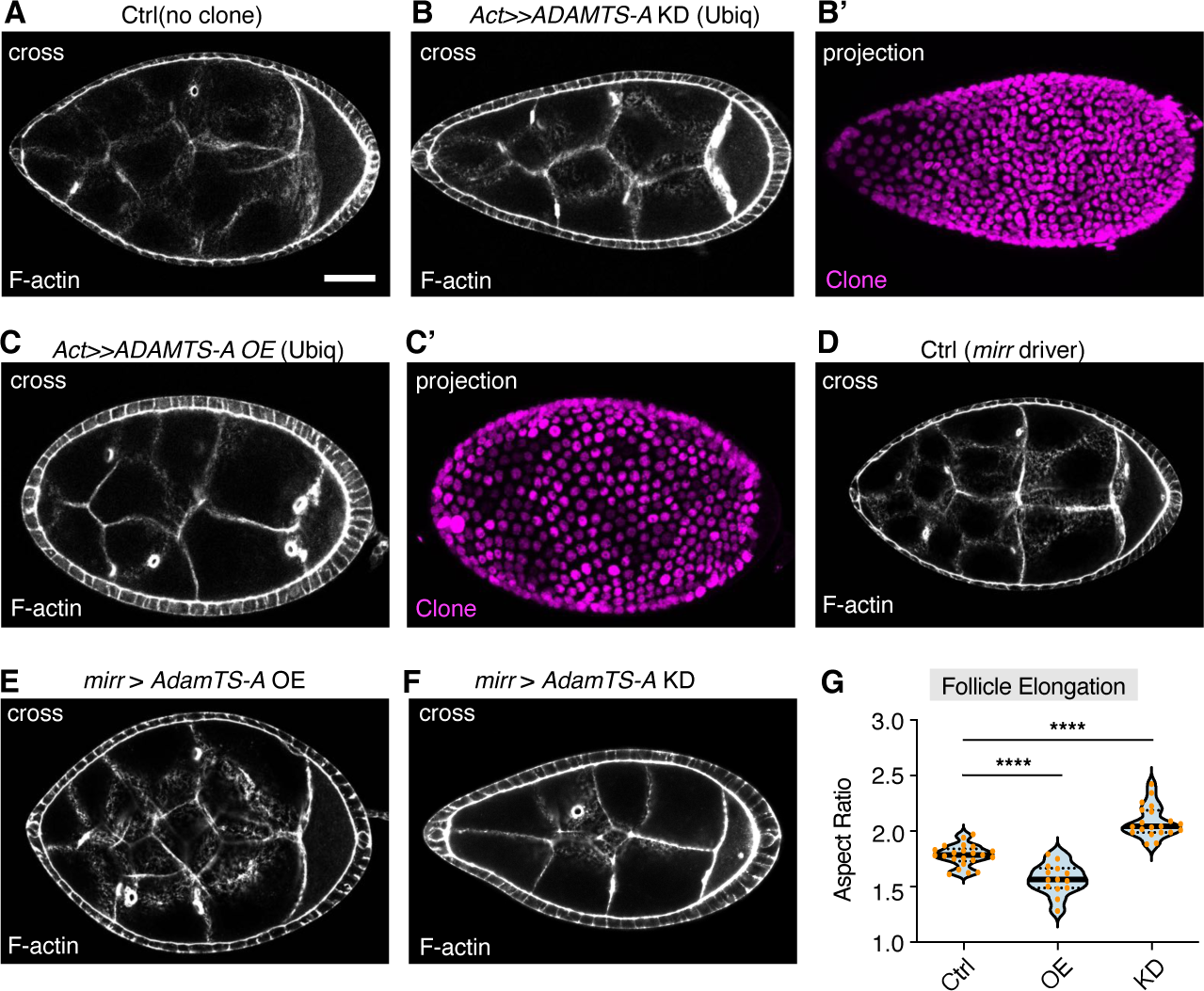
Central ADAMTS-A controls follicle elongation. Compared to control (**A**), ubiquitous depletion (**B**) or elevation of ADAMTS-A (**C**) leads to opposite effects on follicle elongation. Compared to control (**D**), flattening the ADAMTS-A gradient by overexpression in the follicle center via *mirror-GAL4* inhibits tissue elongation (**E**), while steepening the gradient by central depletion enhances elongation (**F**). (**G**) Follicle elongation quantitation by aspect ratio of *mirr*-driven manipulations in D-F. Violin plot with all data points is shown, quartiles and median indicated in dashed or solid line. Statistics: Dunnett’s multiple comparisons, ****P<0.0001. Scale bar: 20 μm.

Shape-determining material properties of follicle tissue can be assessed using an osmotic swelling assay in which living follicles are removed from the animal and placed in water. The resultant influx of fluid creates tissue stresses whose readouts via imaging correlate with BM mechanical values measured by indentation atomic force microscopy (AFM) (17, 18). Two informative parameters are the time and position where the BM breaches as the follicle bursts (**Fig. S1D-F**). WT follicles expand for several minutes before bursting primarily from poles, the areas where BM stiffness is softest. When follicles were depleted of ADAMTS-A with ubiquitous FLP-out, they burst quickly from polar regions. The same phenotype was seen when follicles expressed uniformly high ADAMTS-A levels. Follicles where ADAMTS-A was depleted from the follicle center also burst quickly from polar regions; this phenotype has previously been seen in manipulations that enhance central stiffness. By contrast, overexpression of ADAMTS-A at the center caused rapid bursting preferentially from the center. Overall, either increasing or decreasing ADAMTS-A expression locally or globally disrupts organ shape. The data suggest a requirement for patterned levels of ADAMTS-A.

### ADAMTS-A negatively regulates Col4 levels in the central follicle

Because the A-P pattern of Col4 correlates with BM mechanical properties measured by AFM and bursting as well as organ shape, we examined levels of a functional Col4a2 chain that is endogenously fused with GFP. Interestingly, follicle-wide depletion of ADAMTS-A via ubiquitous FLP-out led to a strong elevation of Col4, but only in the central follicle (**Fig. 3A-B, J**); the terminal regions where *ADAMTS-A* expression is highest showed no evident changes. When ADAMTS-A was overexpressed uniformly, it caused a clear reduction of Col4 in the follicle center (**Fig. 3E, J**), again with no effect evident on the termini. Manipulating ADAMTS-A levels specifically in the follicle center gave similar results: depletion elevated Col4, while overexpression reduced it (**Fig. 3G-I, K**). Analysis of follicle BM deposition with genetic mosaics is challenging because organ rotation leads all cells around the follicle circumference at a particular AP position to contribute to that site (19, 30, 31). However, mosaics that overexpressed ADAMTS-A around an entire circumference site showed elevated Col4 at central but not terminal regions, and could induce localized shape distortions (**Fig. 3C**). By contrast, mosaics that depleted ADAMTS-A only from terminal regions showed no phenotype (**Fig. 3D, J**). These experiments point to an important role for ADAMTS-A in regulating Col4 levels in the follicle center, while the higher levels of ADAMTS-A in termini appear functionally dispensable.

**Figure 3.**
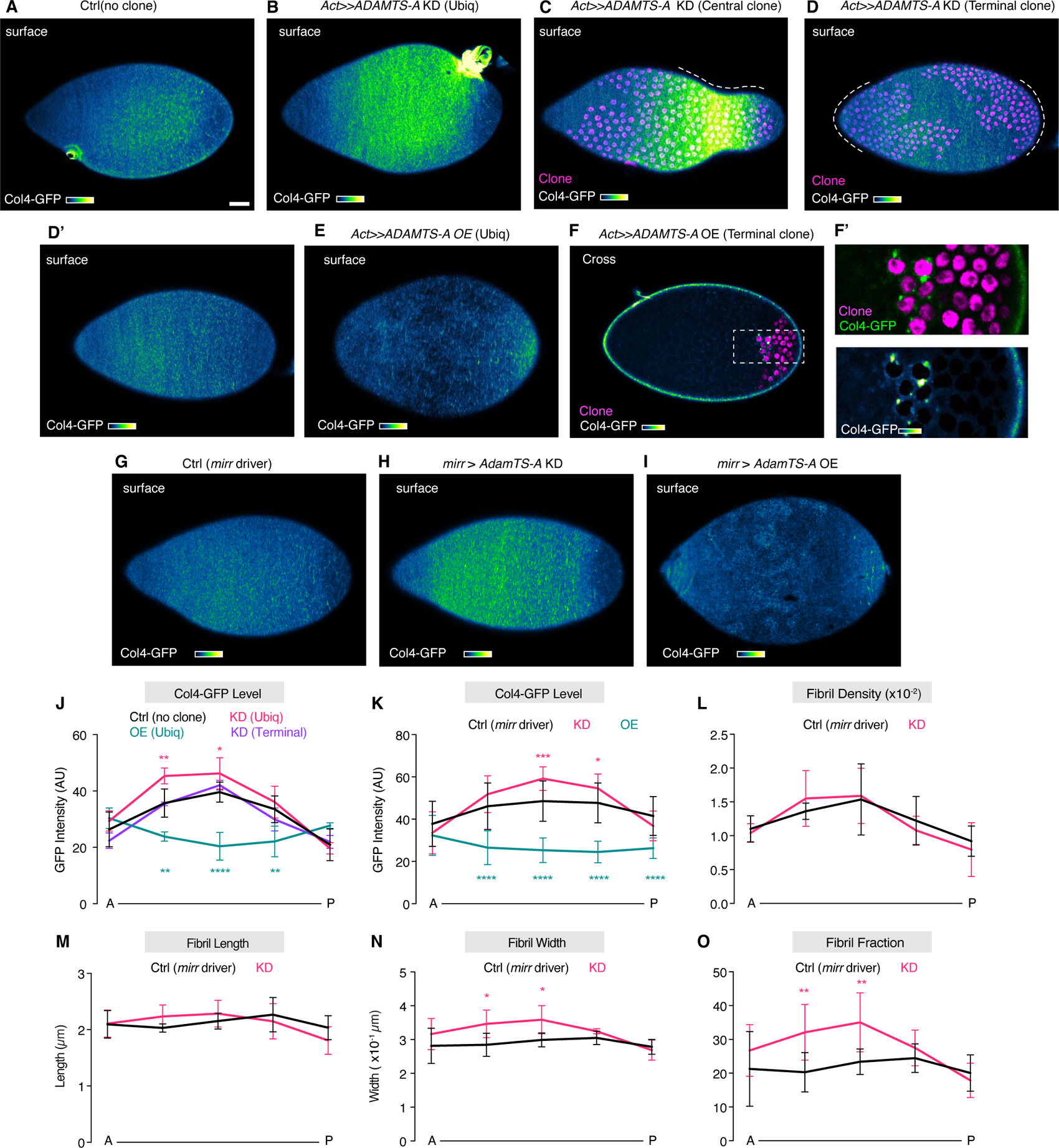
ADAMTS-A attenuates Col4 deposition in the central follicle. Compared to Col4-GFP pattern in control (**A**), follicles ubiquitously depleting ADAMTS*-*A via RNAi (**B**) show elevation in central but not terminal areas. When mosaic clones populate the follicle circumference, knockdown in the central region also shows Col4 elevation (**C**), while no Col4 phenotype is seen with knockdown in the terminus (**D**). Dotted lines in C and D demark regions where mosaic clone is circumferential. Follicles ubiquitously overexpressing ADAMTS-A (**E**) show Col4 depletion in central but not terminal areas; intracellular Col4 trapping in mosaic overexpression clones does not occur at termini (**F**). Col4-GFP patterns when central follicle drivers express control construct (**G**), ADAMTS-A KD (**H**), or ADAMTS-A OE (**I**) show phenotypes similar to ubiquitous expression. (**J**) Quantitates A, B, D and E; (**K**) quantitates G-I, (**L-O**) quantitate Col4-GFP texture: fibril density, length, width, and fibril fraction. Statistics: 2way ANOVA. *P<0.05, **P<0.001, ****P<0.0001. Scale bar: 20 μm.

Col4 in the follicle can be detected in two species: uniform planar expression thought to reflect a typical BM, and dense linear fibrils (19–21). The ratio between the two forms (‘fibril fraction’) is actively regulated. In ADAMTS-A overexpressing follicles, Col4 levels were depleted such that quantitative analysis was not possible. We analyzed follicles in which central depletion of ADAMTS-A causes elevated total Col4 levels and found no changes in fibril length or density, with only modest effects on fibril width (**Fig. 3K-N**). However, fibril fraction was significantly increased (**Fig. 3O**). This phenotype is distinct from known manipulations that increase fibril fraction by elevating Rab10 activity; these favor the generation of longer fibrils and do not alter overall Col4 levels (20). The results suggest that ADAMTS-A’s primary function is to regulate total Col4 deposition, and can also effect its proportional segregation between BM forms.

### ADAMTS-A limits Col4 trafficking to the BM

In follicles overexpressing ADAMTS-A, we noticed a striking phenotype: cells showed pronounced intracellular puncta of Col4 (**Fig. 4 A, B)**. This phenotype is fully penetrant in cells of the central follicle, but nearly absent in cells at terminal regions (**Fig. 3F**). Intracellular Col4 puncta colocalized with or adjacent to markers of ER and the trans Golgi network (**Fig. 4A, B**). Col4 trapping in the ER has been documented when Plod and PH4a, enzymes that hydroxylate Col4 chains to regulate dimer- and trimerization, are depleted (32, 33). However, upon ADAMTS-A overexpression Col4 was trapped in multiple sites distributed throughout the follicle cell, distinct from the basal sites seen in PH4a-depleted cells (**Fig. 4C-E**). Although reagents to detect endogenous ADAMTS-A are not available, we analyzed GAL4-driven ADAMTS-A-GFP and found extensive intracellular localization that partially overlapped with markers of the ER, as also seen in the salivary gland, along with expression at the basal cell surface (34) (**Fig. 4F**). Together, these data point to an undocumented relationship between ADAMTS-A and the Col4 secretory pathway.

**Figure 4.**
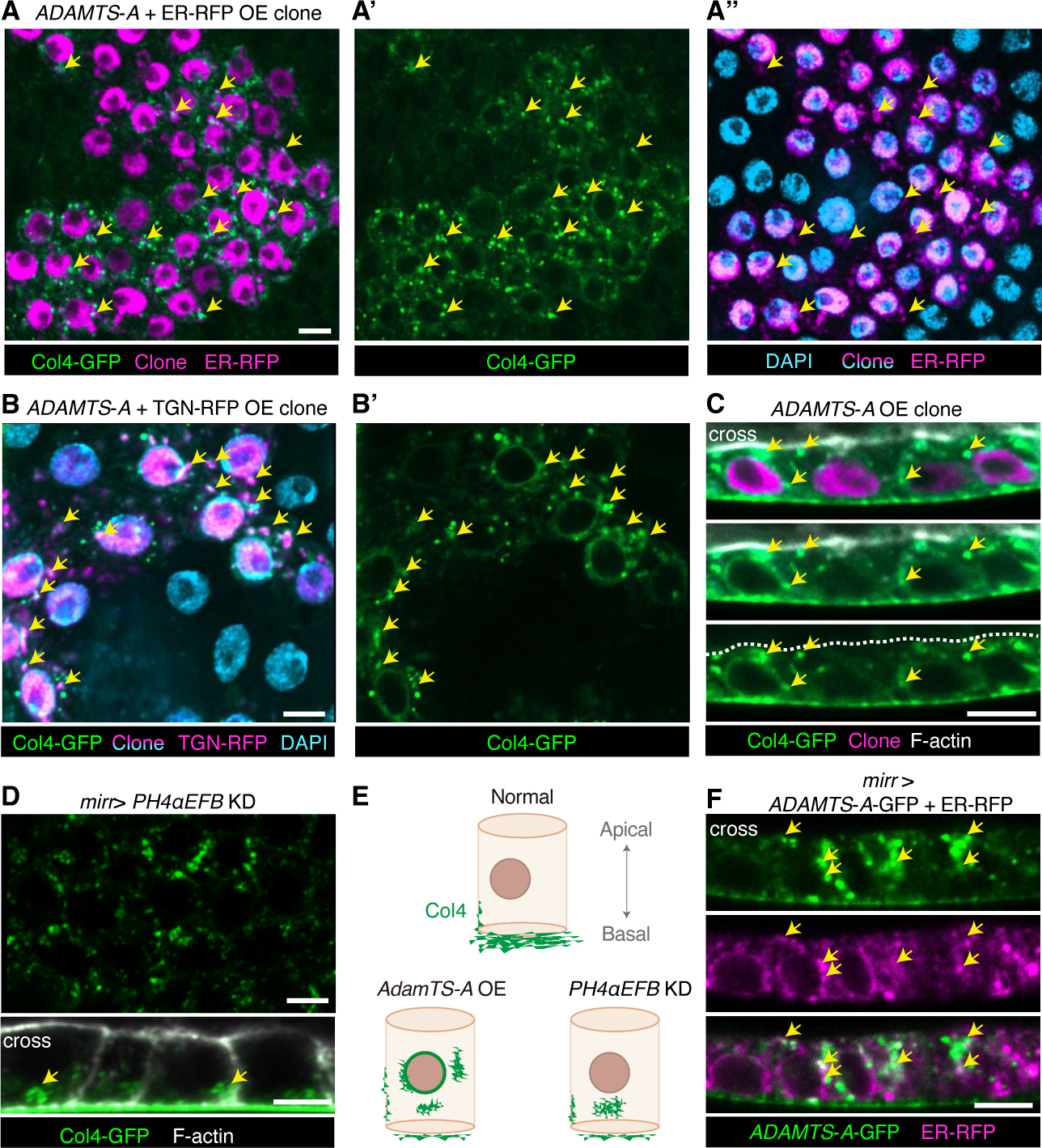
ADAMTS-A regulates Col4 secretion. **(A)** Sub-basal section of central mosaic clone overexpressing ADAMTS-A along with the ER marker KDEL-RFP. Unlike neighboring control cells, ADAMTS-A overexpressing cells show Col4-GFP intracellular punctate that colocalize with or adjacent to the ER (arrows). **(B)** Intracellular Col4-GFP colocalizes with TGN-RFP (trans-Golgi, arrows) in central clones overexpressing ADAMTS-A. (**C**) Cross-sections of ADAMTS-A overexpressing cells show intracellular Col4-GFP (arrows) scattered along the apical-basal axis. (**D**) Depletion of PH4αEFB leads to Col4-GFP punctate restricted near basal surface (arrows). (**E**) Schematic representations of normal, ADAMTS-A overexpressing, and PH4αEFB KD cells. Col4-GFP is normally predominantly located in the BM, but elevated ADAMTS-A causes Col4 trapping along the perinuclear, ER, and TGN; PH4αEFB depletion leads to basal intracellular punctate. (**F**) ADAMTS-A-GFP driven in central follicle overlaps with or is adjacent to ER (arrows).

### ADAMTS-A inhibits Rab-mediated secretion and deposition of Col4

To analyze dynamics of Col4 within the BM, we used CRISPR to tag Col4a2 with the photoconvertible protein mMaple. Like animals carrying Col4a2-GFP, animals carrying Col4a2-mMaple are homozygous viable with no evident defects, indicating negligible impact on Col4 function. We used irradiation to switch mMaple fluorescence from green to red spectrum, allowing inference of two processes (**Fig. 5A**). Fluorescence recovery of green signal (FRAP) is consistent with Col4 molecules that are newly deposited into the BM, while fluorescence decay of red signal (FDAP) can reveal the half-life of Col4 previously deposited into the BM. Interpretation of these data requires caveats but is informed by the fact that ∼80% of Col4 at st. 7-8 is produced by follicle epithelial cells rather than recruited from extrinsic sources (35, 36). Strikingly, ADAMTS-A depletion in follicles led to faster FRAP and an increased mobile fraction (**Fig. 5B-D**), suggesting enhanced deposition into the BM. By contrast, depletion of ADAMTS-A did not alter Col4a2-mMaple FDAP, implying that this metalloprotease does not influence follicle BM degradation (**Fig. 5E**).

**Figure 5.**
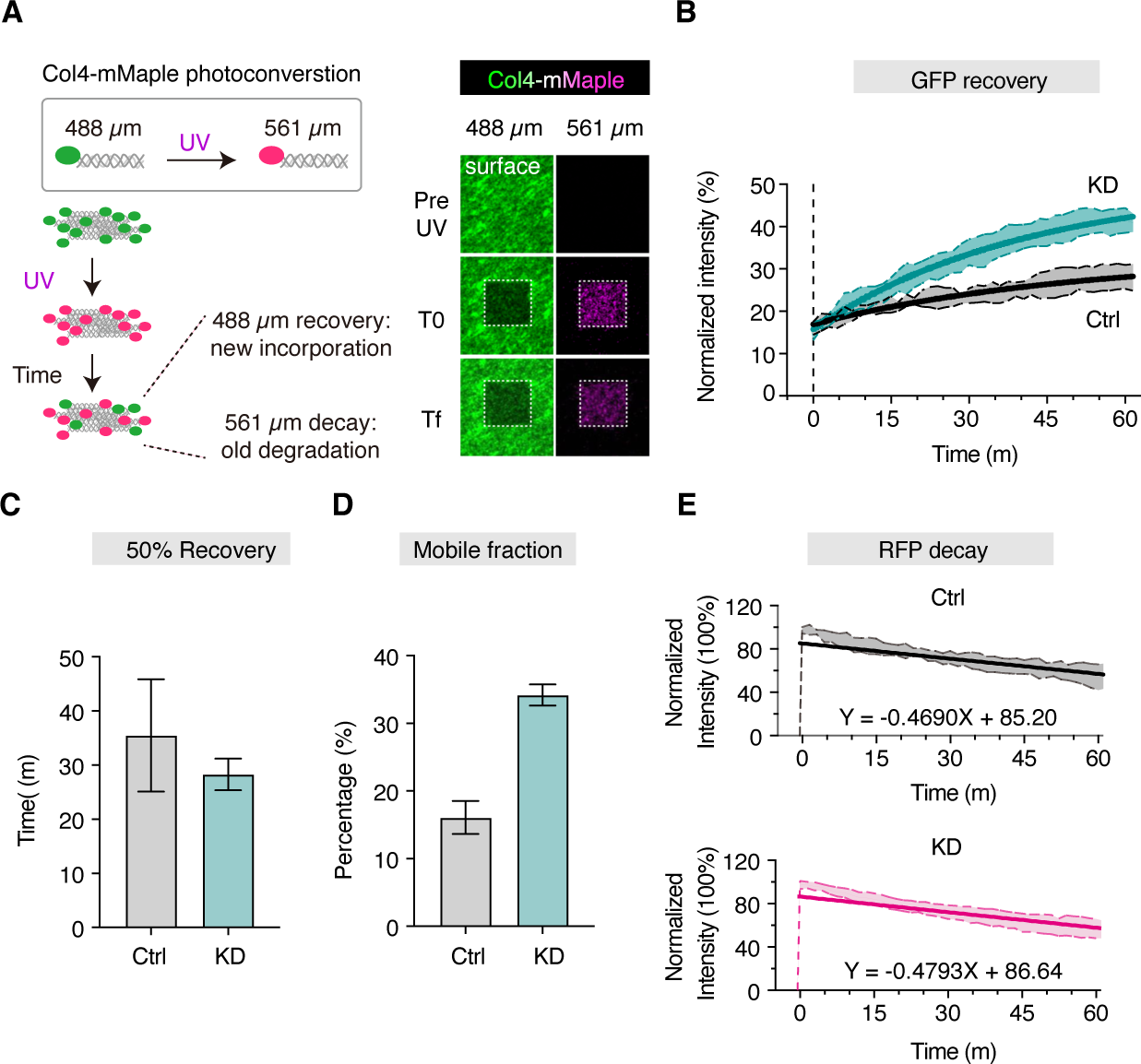
ADAMTS-A restrains Col4 incorporation without modulating extracellular dynamics. (**A**) Schematic of Col4-mMaple photoconversion. Upon UV irradiation (T0), mMaple fluorescence shifts from green to red spectrum. In BM, decline of converted Col4-mMaple in red at Tf represents Col4 decay, whereas recovery in green reflects newly deposited Col4. (**B-D**) FRAP assay of Col4-mMaple in green spectrum of control and ADAMTS-A KD follicles. Fitted non-linear curve ± SD is plotted. Depletion of ADAMTS-A leads to faster recovery and increased mobile fraction of Col4-mMaple, as quantitated in C and D (mean and SEM). (**E**) Decay of Col4-mMaple in red spectrum. Equations from simple linear regression of control and ADAMTS-A KD mutant indicated at graph bottom. Fitted lines ± SD shown. Half-life is 90.8 and 90.4 minutes in control and ADAMTS-A KD, respectively.

**Figure 6.**
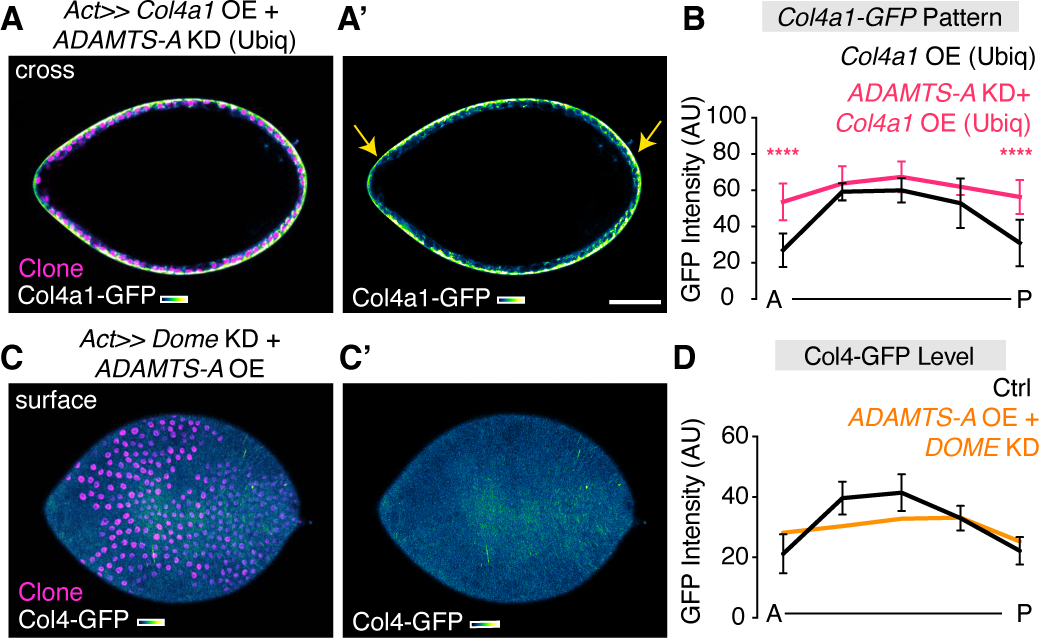
Terminal ADAMTS-A limits excess Col4 production. While uniform overexpression of *Col4a1-GFP* transcript in WT follicles leads to strongly reduced Col4a1-GFP protein incorporation into terminal BM (see Fig. 1F), co-depletion of ADAMTS-A allows uniform incorporation along the AP axis, including in termini (arrows, **A**, quantitated in **B**). When Col4 levels should be uniformly high due to blockage of the terminal patterning signal via Dome depletion, restoring ADAMTS-A via overexpression maintains low terminal Col4 levels (**C**, quantitated in **D**).

Through what mechanism might ADAMTS-A impact Col4 deposition? Col4 secretion from follicle epithelial cells to the BM involves specialized trafficking routes that are regulated by Rab proteins. Although details are not fully understood, Rab10 promotes fibril formation, while both Rab10 and Rab8 can support trafficking of Col4 to the basal cell surface and incorporation into the planar matrix (33, 37, 38). Examining endogenously fluorescent-tagged functional versions of Rab proteins, we found that ADAMTS-A overexpression caused a ∼50% decrease in levels of Rab10, as well as Rab8 and Rab11 **(Fig. S2A-F)**. The ADAMTS-A overexpression phenotype, where Col4 remains trapped within the cell, resembles that documented when all three Rab proteins are depleted (37). By contrast, ADAMTS-A depletion caused a mild increase in Rab10 levels but not Rab8 or Rab11 **(Fig. S2A-F)**. Interestingly, when the Rab10 effector EHBP1 was coexpressed with ADAMTS-A, Col4 intracellular trapping was resolved and fibril formation was restored **(Fig. S2G).** Taken together, the data are consistent with a model in which ADAMTS-A influences Rab10-dependent exit of Col4 from the secretory pathway to limit its deposition into the BM.

### A role for terminal ADAMTS-A in the Col4 gradient

The results above indicate that ADAMTS-A can control levels of Col4 incorporation in the central follicle. A major feature of the Col4 gradient is its low point at each follicle terminus, which depends on Upd secreted from polar cells to activate STAT in nearby terminal epithelial cells (17, 23). Both ubiquitous and mosaic depletion studies above show that the high terminal levels of ADAMTS-A, a known transcriptional target of STAT (26), were not necessary to maintain low terminal Col4 levels (**Figs. 3A-C, J**). However, ADAMTS-A depletion was sufficient to allow incorporation of overexpressed *Col4a1* into the BM at follicle termini (**Fig. 5A**), in contrast to the absence of incorporation when ADAMTS-A is present (**Fig. 1F, Fig. 5B**). Moreover, the elevated terminal Col4 seen when STAT signaling is blocked could be inhibited by restoring ADAMTS-A expression (17) (**Fig. 5C-D**). These results are consistent with a model in which Upd ensures the organ-shaping gradient by transcriptionally regulating *ADAMTS-A* to limit incorporation of excess Col4 in termini.

## Discussion

Recent studies have revealed numerous instances across animal phylogeny in which ECM is created, modified or destroyed at specific sites to direct morphogenesis (3). How do discrete cells impose a spatial pattern upon the ECM’s continuous extracellular material? Here we explore one such pattern, the symmetric low-high-low gradient of Col4 within the fly follicle BM. Rather than simple control of Col4 transcription by the polar cell morphogen Upd, we find unexpected complexity that involves regulation of exocytic exit by regional levels of ADAMTS-A. The intracellular activity proposed here for the ADAMTS-A metalloprotease alters total Col4 deposition independently of the extracellular activity of the secreted matrix metalloprotease MMP1 that regulates Col4 fibril texture (13). Layered cell-autonomous as well as non-autonomous modes of Col4 regulation may arise from the necessity of creating fine pattern and incorporating dynamic feedback to reach precise follicle shape, and provide a rationale for why this organ synthesizes its own BM rather than assembling it solely from soluble sources.

We provide evidence that ADAMTS-A limits follicle Col4 levels through a mechanism involving the secretory pathway. ADAMTS-A and its orthologs are documented regulators of ECM, but they do not appear to have a single simple molecular mechanism of doing so. Amongst the 19 human ADAMTS family members, the closest orthologs are ADAMTS9 and ADAMTS20, originally suggested to cleave proteoglycans in extracellular space. Yet ADAMTS-A shares a conserved C-terminus with ADAMTS9 and the *C. elegans* ortholog GON-1, which also localize to and promote ER to Golgi secretory transport (39, 40). ADAMTS-A’s role in Col4 trafficking appears distinct from that described for GON-1 and ADAMTS9, in that it negatively regulates exit from the secretory pathway, but our data nevertheless extend the evidence that these proteins have intracellular as well as extracellular roles.

While Col4 is unlikely to be a direct substrate, ADAMTS-A is a potent negative regulator of Col4 levels, shown here for the central follicle and previously for the larval glia, where it also inhibits tissue stiffening (41). GON-1 negatively regulates Col4 in the worm gonad, but can also positively promote Col4 function in the linkage between adjacent BMs; in presynaptic boutons both Col4-promoting and -antagonizing roles have been proposed (42–45). In the embryonic salivary gland, ADAMTS-A releases migrating cells from the apical ECM through a process in which Col4 is not implicated (40). Finally, ADAMTS-9 is mutated in human patients with fibrotic nephropathies that could involve an intracellular mode of action (46). The identification of bona fide *in vivo* substrates of ADAMTS-A and its orthologs, which has yet to be achieved, will shed important light on the molecular mechanisms by which these proteins control morphogenesis.

The growing follicle faces the challenge of achieving a precise morphogenetic outcome from a quite limited set of initial patterning information. Early follicle epithelial cells are homogenous; the only cue that diversifies their fates is the morphogen Upd, secreted from the two pairs of PCs that are incorporated into the epithelium from a distinct cell lineage (22, 47). PCs are thus responsible not only for distinguishing central vs terminal cell fates to create different eggshell structures and convey pattern to the embryo, but also for distinguishing central from terminal BM mechanical properties to allow a specific degree of tissue elongation. PCs organize morphogenesis by three distinct mechanisms: creation of an ADAMTS-A gradient that regulates graded levels of Col4 incorporation, secretion of the matrix metalloprotease MMP1 to provide more sensitive tuning of fibril ‘texture’, and regulation of terminal cell behaviors including contractile apical pulses (13, 23). These functions are not sufficient to account for all BM patterning; for instance the direct effectors that establish lower terminal Col4 levels compared to the center remain to be discovered, as do mechanisms that set the distributions of Laminin and Perlecan which are distinct from that of Col4 (17, 18, 48). Further analysis of this model system is likely to continue to yield unexpected mechanisms by which the abundant extracellular environment of animal bodies is patterned.

## MATERIALS AND METHODS

### Fly Strains and Husbandry

Stocks and genotypes used in this study are described is **Supplemental Tables 1 and 2**. Cornmeal molasses food with dried yeast powder were used to rear fly stocks. Experimental crosses were raised at 25°C unless indicated otherwise. For temperature shift experiments (*GAL4*/*GAL80*^TS^), animals were kept at 25°C before third instar (L3), then shifted to 18°C before eclosion. Adults eclosed within 2 days were collected and shifted to 30°C for 2 days before dissection. FLPout clonal inductions were carried out by heat-shocking L3 animals at 37°C for 13 minutes, then returning to 18°C until eclosion. Animals were transferred to 30°C for 2 days prior to dissection.

### Generation of Col4-mMaple

CRISPR-Cas9-mediated homologous recombination was carried out to generate the Col4-mMaple fly line. mMaple coding sequence was inserted at the same site as the GFP in *vkg^CC00791^*-*GFP*. GFP positive fly line with strong and stable basement membrane Col4-mMaple signal was used for photoconversion.

### Immunofluorescent Staining and Imaging

Ovaries were dissected in Schneider’s medium before 4% paraformaldehyde (PFA) fixation for 15 minutes. After serial PBS washes, samples were stained with Phalloidin and/or DAPI before being mounted in antifade (Diamond Antifade Mountant, Invitrogen). GFP signals were boosted using nanobody staining (1:1000, ChromoTek) for Rab8-YFP, Rab10-YFP, or Rab11-YFP. Image acquisitions using Zeiss LSM 700 or 900 microscopes were conducted within 48 hours post sample preparations. Objectives used were 20x Plan-Apochromat/0.8 or 40x LD C-Apochromat/NA 1.1 W Korr. Pinhole settings were 1 Airy unit for all experiments.

### In Situ Hybridization Chain Reaction (HCR)

Probe sets for target genes (*ADAMTS*-*A* and *vkg/Col4a2*), amplifier hairpins, and buffers were obtained from Molecular Instruments. Conserved sequences in alternative transcripts were used as targeting regions to generate probe sets. Reactions was carried out based on HCR v3.0. protocol (49). Briefly, muscle-free ovarioles were dissected in culture medium and subjected to 4% PFA fixation for 15 minutes. After serial washes, samples were permeabilized by detergent solution for 30 minutes. Ovarioles were then incubated in 10 nM of probe sets in hybridization solution at 37°C overnight. Following serial washes, samples were incubated in fluorophore-tagged hairpin amplifiers (H1 and H2: 90 nM each) overnight at room temperature. DAPI counter-staining was performed prior to mounting for stage determination. Negative control was carried in follicles subjected only to amplifiers without probe sets.

### Photoconversion of Col4-mMaple

*Ex vivo* cultures of stage 8 Col4-mMaple follicles were performed as described (19) with osmolality of culture medium calibrated to 260 Osm/L. Images were acquired in dual-channels for both pre- and post-405 nm laser photoconversions. Z stacks of 7 μm (0.5 μm spacing) were acquired every 30 seconds (pre) or 90 seconds (post) for 2 minutes or 1 hour, respectively, through a 40x objective with 2-fold digital zoom, in 512 x 512-pixel resolution (dwell time: 1.52 μs, total area as 79.9 x 79.9 μm^2^). Photoconversions were carried out in an 8 x 8 μm^2^ region, as the 405 nm laser power turned to 50% for 10 iterations (dwell time: 1.52 μs). Differential z-position during photoconversion was used to focus on the BM plane for maximal efficiency. Software autofocus was carried out referencing absolute distance from cover glass. Focus strategy was defined for Z values/Focus surface per sample with local adaptive Z that updates with single offset and standard stabilization. Image registrations were carried out in ZenBlue through embedded Time Alignment modules with translocation and linear interpolation. GFP spectrum recovery (FRAP) was quantified as previously described (50). Normalized decay in RFP spectrum over time (I_t_) was quantified using I_t_= (i_t_-i_min_) /(i_max_-i_min_), where the i_t_, i_max_, and i_min_ stands for the raw, maximum, and minimum pixel intensity. Raw pixel values and image processes were retrieved through FIJI; normalization, statistics, and graph plotting were performed in Prism10.

### Data Quantification and Statistics

For follicle elongation, bursting assay, Col4-GFP intensity and fibril analysis were performed as described previous (13). 3D projected images of HCR experiments were used to measure organ-level signal intensity of developing follicle. Single-cell RNA sequence dataset was obtained from Mendeley Data (47). *Col4a1*, *Col4a2*, and *ADAMTS-A* in early-to-mid follicle clusters were retrieved and plotted using Seurat 5. Statistical analysis used in experiments were: One -way ANOVA: follicle elongation, burst time, and Rab10/8/11-YFP intensity; Two-way ANOVA: A-P GFP intensity, fibril density, fibril length, fibril weight, fibril fraction, and secreted-GFP; Chi-squared using Bonferroni correction: burst position.

## ACKNOWLEDGMENTS

We thank Laura Mathies for generating Vkg-mMaple, the Hariharan lab for advice on HCR, and the Bilder lab for helpful discussion. We acknowledge generous gifts of reagents from Afshan Ismat, Debbie Andrew, Trudi Schupbach and the community resources provided by TRiP at Harvard Medical School (NIH/NIGMS R01-GM084947), the Bloomington Drosophila Stock Center (NIH P40OD018537) and the Vienna Drosophila Resource Center. This work was supported by NIH grant GM130388 to D.B.

