## Supplemental materials for "Basement membrane patterning by spatial deployment of a secretion-regulating protease"

### Supplemental Figure Legends

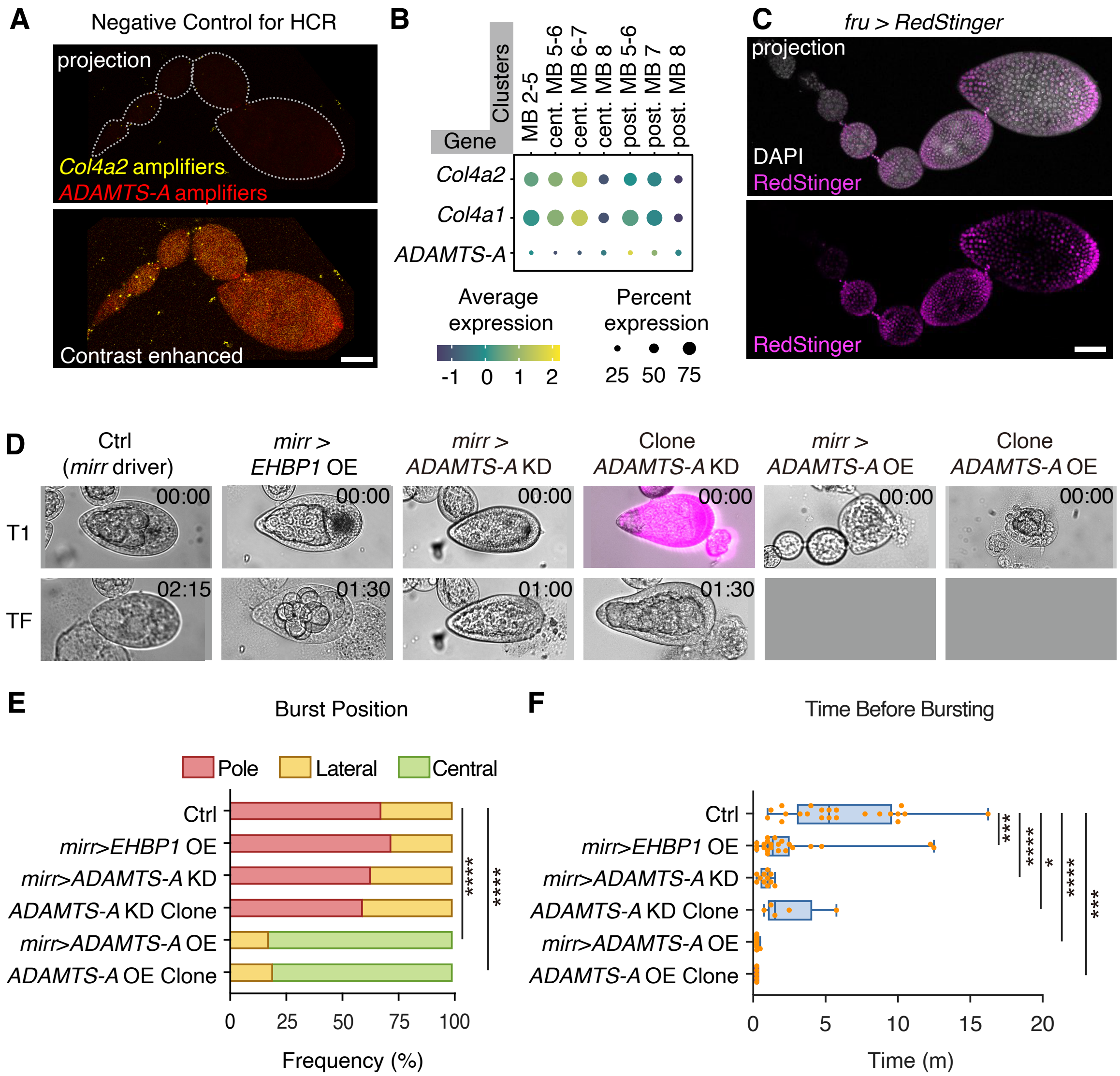


**Figure S1. ADAMTS-A patterning regulates tissue mechanics.**
(A) Negative control for HCR detection of Fig. 1D: follicles incubated with fluorophore amplifiers but not target RNA probe sets. Upper image shows same imaging conditions as Figure 1D; lower image shows contrast enhanced. (B) Multi-group dotplot of *Col4a1, Col4a2* and *ADAMTS-A*: single-cell RNAseq of follicle clusters stages 2-8 retrieved from Rust et al. 2020. (C) *Fruitless-GAL4* driver (magenta) in developing follicles.(D) Bursting assay for control (*mirr* driver), central stiffening (*mirr>EHBP1*), central or ubiquitous ADAMTS-A depletion (*mirr> ADAMTS-A KD* or *ubi FLPout>ADAMTS-A KD*, respectively), and central or ubiquitous ADAMTS-A overexpressing follicles (*mirr>ADAMTS-A OE* or *ubi FLPout>ADAMTS-A OE*, respectively). Upper images: first time-point upon osmotic challenge. Lower images: time points when follicles burst due to osmotic swelling. Time stamps in upper right reflect minutes:seconds. (E) Quantitation of spatial burst position from D. Statistics: Chi-squared test, Bonferroni correction for 6 groups; ****P<0.0001. (F) Quantitation of burst time from D. *Statistics:* Dunnett's multiple comparisons, *P<0.05, ***P<0.001, ****P<0.0001. Scale bar: 20 μm.

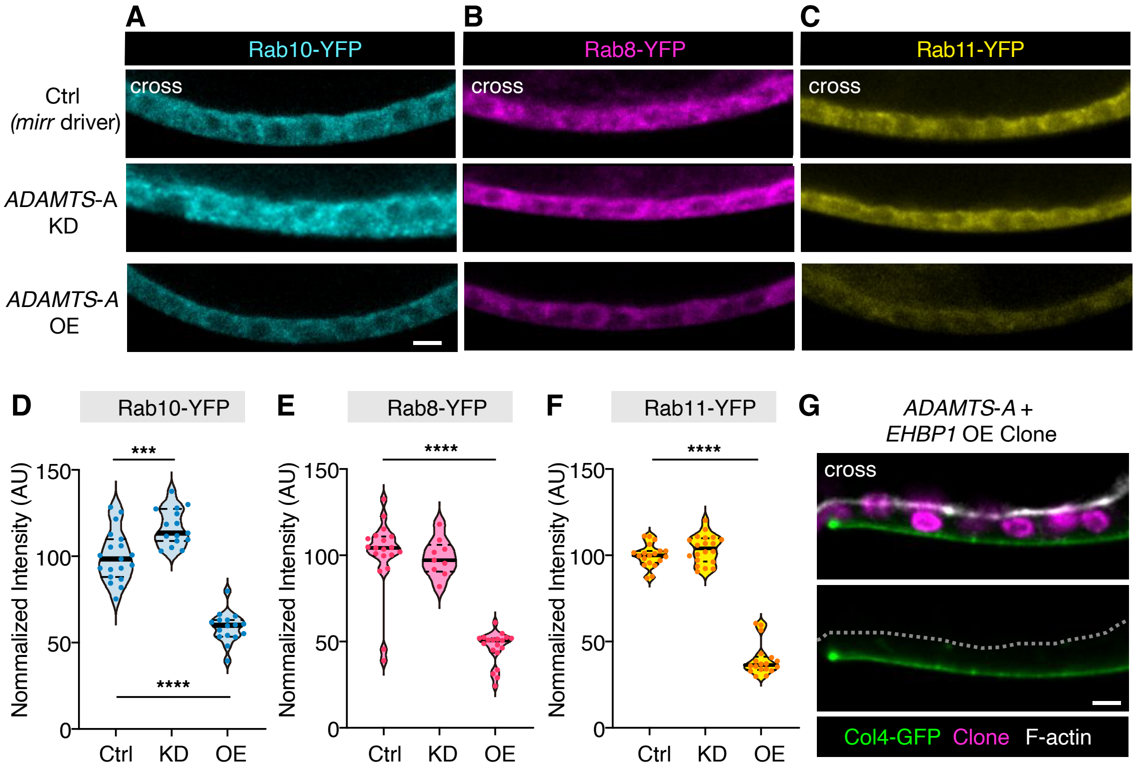


**Figure S2. ADAMTS-A modulates Rab protein levels**

(A-C) Endogenously tagged Rab proteins in central epithelial sections of follicles expressing *mirr-GAL4*. (A) shows Rab 10-YFP in control (top), ADAMTS-A KD (middle) or ADAMTS-A overexpressing follicles. (B) and (C) show Rab8-YFP and Rab11-YFP (D) in control (mirr driver), ADAMTS-A KD, or ADAMTS-A OE follicles. Scale bar: 20 μm. Quantitated in (D-F). Statistics: Tukey's multiple comparisons. ***P<0.0005, ****P<0.0001. (G) EHBP1 overexpression resolves the Col4 trapping caused by ADAMTS-A overexpression; compare Figs. 3F and 4A-C. *Scale* bar: 5 μm.

**Supplementary Table 1. Genotypes, experimental conditions, and sample size**

| | | | | |
|---|---|---|---|---|
| Figure 1 | B | *vkg-GFP* (II) | 25 °C; 3 D | n= 8 |
| | C | *10xSTAT92E-GFP* (III) | 25 °C; 3 D | n= 12 |
| | D | *w*$^{1118}$ | 25 °C; 3 D | n= 13 |
| | E | *w*$^{1118}$ | 25 °C; 3 D | n= 7 (st3-4);<br>8 (st5-6);<br>7 (st7-8) |
| | F | *hs-FLP*$^{122}$*; Sp/UAS-cg25c-GFP; Act>STOP>GAL4,UAS-His-RFP/ +* | 30 °C; 3 D | n= 9 |
| Figure 2 | A | *hs-FLP*$^{122}$*; + ; Act>STOP>GAL4,UAS-His-RFP/ +* | 30 °C; 2 D | n=4 |
| | B | *hs-FLP*$^{122}$*; UAS-ADAMTS-A KD/+;<br>Act>STOP>GAL4,UAS-His-RFP/ +* | 30 °C; 2 D | n=5 |
| | C | *hs-FLP*$^{122}$*; UAS-ADAMTS-A OE/+;<br>Act>STOP>GAL4,UAS-His-RFP/ +* | 30 °C; 2 D | n=4 |
| | D | *tub-GAL80*$^{TS}$*/+; mirr-GAL4/+* | 30 °C; 2 D | n= 22 |
| | E | *tub-GAL80*$^{TS}$*/UAS-ADAMTS-A OE; mirr-GAL4/+* | 30 °C; 2 D | n= 14 |
| | F | *tub-GAL80*$^{TS}$*/UAS-ADAMTS-A KD; mirr-GAL4/+* | 30 °C; 2 D | n= 20 |
| | G | as D-F | | |
| Figure 3 | A | *hs-FLP*$^{122}$*; vkg-GFP/+ ; Act>STOP>GAL4,UAS-His-RFP/ +* | 30 °C; 2 D | n= 5 |
| | B-D | *hs-FLP*$^{122}$*; vkg-GFP/UAS-ADAMTS-A KD ;<br>Act>STOP>GAL4,UAS-His-RFP/ +* | 30 °C; 2 D | n= 5(Ubi);<br>2(Cen);<br>4(Ter) |
| | E-F | *hs-FLP*$^{122}$*; vkg-GFP/+ ; Act>STOP>GAL4,UAS-His-RFP/ UAS-ADAMTS-A OE* | 30 °C; 2 D | n= 3(Ubi);<br>4 (Ter) |
| | G | *tub-GAL80*$^{TS}$*/ vkg-GFP; mirr-GAL4/+* | 30 °C; 2 D | n= 22 |
| | H | *tub-GAL80*$^{TS}$*/ vkg-GFP; mirr-GAL4/UAS-ADAMTS-A KD* | 30 °C; 2 D | n= 21 |
| | I | *tub-GAL80*$^{TS}$*/ vkg-GFP; mirr-GAL4/UAS-ADAMTS-A OE* | 30 °C; 2 D | n= 14 |
| | J | as G-I | | |
| | K-N | as G-H | | |
| Figure 4 | A | *hs-FLP*$^{122}$*; vkg-GFP/UAS-ADAMTS-A OE ;<br>Act>STOP>GAL4,UAS-His-RFP/UAS-KDEL-RFP* | 30 °C; 2 D | n= 13 |
| | B | *hs-FLP*$^{122}$*; vkg-GFP/UAS-GalT-TagRFP ;<br>Act>STOP>GAL4,UAS-His-RFP/ UAS-ADAMTS-A OE* | 30 °C; 2 D | n= 9 |
| | C | *hs-FLP*$^{122}$*; vkg-GFP/+ ;<br>Act>STOP>GAL4,UAS-His-RFP/ UAS-ADAMTS-A OE* | 30 °C; 2 D | n= 7 |

| | | | | |
|---|---|---|---|---|
| | D | *tub-GAL80$^{TS}$/ vkg-GFP; mirr-GAL4/ UAS-PH4αEFB KD* | 30 °C; 3 D | n= 10 |
| | F | *tub-GAL80$^{TS}$/ UAS-ADAMTS-A-GFP ; mirr-GAL4/ UAS-KDEL-RFP* | 30 °C; 2 D | n= 7 |
| Figure 5 | A | *vkg-mMaple* (II) | 25 °C; 2 D | n= 6 |
| | B-D | *tub-GAL80$^{TS}$/ vkg-mMaple; mirr-GAL4/+* | 30 °C; 2 D | n= 5 |
| | | *tub-GAL80$^{TS}$/ vkg-mMaple; mirr-GAL4/ UAS-ADAMTS-A KD* | 30 °C; 2 D | n= 8 |
| Figure 6 | A | *hs-FLP$^{122}$; Sp / UAS-cg25c-GFP; Act>STOP>GAL4,UAS-His-RFP/ UAS-ADAMTS-A KD* | 30 °C; 2 D | n= 7 |
| | B | *as A, Fig. 1F* | 30 °C; 2 D | |
| | C | *hs-FLP$^{122}$; vkg-GFP / UAS-Dome KD; Act>STOP>GAL4,UAS-His-RFP/ UAS-ADAMTS-A OE* | | n= 1 |
| | D | *as C,* Ctrl: no clonal induction | | n= 3(Ctrl) |
| Sup 1 | A | *w$^{1118}$* | 25 °C; 3 D | n= 8 |
| | C | *tub-GAL80$^{TS}$/ + ; fru-GAL4/ UAS-Red-Stinger* | 30 °C; 2 D | n= 15 |
| | D | *tub-GAL80$^{TS}$/ +; mirr-GAL4/+* | 30 °C; 2 D | n= 22 |
| | | *tub-GAL80$^{TS}$/ +; mirr-GAL4/UAS-FLAG-EHBP1* | 25 °C; 3 D | n= 22 |
| | | *tub-GAL80$^{TS}$/ +; mirr-GAL4/UAS-ADAMTS-A KD* | 30 °C; 2 D | n= 11 |
| | | *hs-FLP$^{122}$; vkg-GFP/+ ; Act>STOP>GAL4,UAS-His-RFP/ UAS-ADAMTS-A KD* | 30 °C; 2 D | n= 5 |
| | | *tub-GAL80$^{TS}$/ +; mirr-GAL4/UAS-ADAMTS-A OE* | 30 °C; 2 D | n= 11 |
| | | *hs-FLP$^{122}$; vkg-GFP/+ ; Act>STOP>GAL4,UAS-His-RFP/ UAS-ADAMTS-A OE* | 30 °C; 2 D | n= 5 |
| | E-F | as D | | |
| Sup 2 | A,D | *Rab10-YFP/+; tub-GAL80TS/ + ; mirr-GAL4/ +* | 30 °C; 2 D | n= 19 |
| | | *Rab10-YFP/+; tub-GAL80TS/ UAS-ADAMTS-A KD ; mirr-GAL4/ +* | 30 °C; 2 D | n= 15 |
| | | *Rab10-YFP/+; tub-GAL80TS/ UAS-ADAMTS-A OE ; mirr-GAL4/ +* | 30 °C; 2 D | n= 16 |
| | B,E | *tub-GAL80$^{TS}$/ +; mirr-GAL4/Rab8-YFP* | 30 °C; 2 D | n= 19 |
| | | *tub-GAL80$^{TS}$/ UAS-ADAMTS-A KD; mirr-GAL4/Rab8-YFP* | 30 °C; 2 D | n= 9 |
| | | *tub-GAL80$^{TS}$/ UAS-ADAMTS-A OE; mirr-GAL4/Rab8-YFP* | 30 °C; 2 D | n= 19 |
| | C,F | *tub-GAL80$^{TS}$/ +; mirr-GAL4/Rab11-YFP* | 30 °C; 2 D | n= 19 |
| | | *tub-GAL80$^{TS}$/ UAS-ADAMTS-A KD; mirr-GAL4/Rab11-YFP* | 30 °C; 2 D | n= 20 |

| | | | | |
|---|---|---|---|---|
| | | *tub-GAL80$^{TS}$/ UAS-ADAMTS-A OE; ßmirr-GAL4/Rab11-YFP* | 30 °C; 2 D | n= 18 |
| | G | hs-FLP$^{122}$; vkg-GFP/UAS-ADAMTS-A OE ; Act>STOP>GAL4,UAS-His-RFP/ UAS-FLAG-EHBP1 | 30 °C; 2 D | n= 18 |

**Supplementary Table 2: Fly Strain, Source, and Identifier**

| | | |
|---|---|---|
| Bloomington Stock Center | *w$^{1118}$* | BL 5905 |
| | *hs-FLP$^{122}$; Act5c>CD2>GAL4, UAS-RFP* | BL 30558 |
| | *Rab8-YFP* | BL 62546 |
| | *Rab10-YFP* | BL 62548 |
| | *Rab11-YFP* | BL 62549 |
| | *UAS-dome*-RNAi | BL 7019 |
| | *UAS-PH4α*-RNAi | BL 33695 |
| | *UAS-RFP-KDEL* | BL 30910 |
| Vienna Drosophila Resource Center | LamininB1-GFP | v318180 |
| | *UAS*-ADAMTS-A RNAi | v33347; v110157 |
| Gifted | *UAS-ADAMTS-A* | Trudi Schupbach; PMID: 30385460 |
| | *UAS-ADAMTS-A* | Afshan Ismat; PMID: 23536567 |
| | UAS-*ADAMTS-A-GFP* | Afshan Ismat; PMID: 23536567 |
| | *UAS-cg25c-GFP* | Stéphane Noselli; PMID: 26456819 |
| | *vkg$^{CC00791}$-GFP* | Allan C. Spradling; PMID:17194782 |
| | *mirr-GAL4* | Anne-Marie Pret; PMID: 19882738 |
| | *fruitless-GAL4* | Anne-Marie Pret; PMID: 23222440 |
